# Effect of different lining paper materials and infusions on oviposition preference of *Aedes aegypti* (Diptera: Culicidae) gravid mosquitoes

**DOI:** 10.1101/2024.09.22.613174

**Authors:** Mahfuza Momen, Kajla Seheli, Md. Aftab Hossain, Ananna Ghosh, Md. Forhad Hossain

## Abstract

*Aedes aegypti* (Linnaeus) mosquito is a vector, responsible for increasing global public health concerns due to its rapid geographical spread and increasing vectorial capacity. Understanding the breeding behavior of *Ae. aegypti* is crucial to control this vector and thereby limiting the spread of diseases. To study this mosquito breeding in the laboratories and ovitraps, different types of lining-papers, surrounding inner surface of a container along with different types of added infusions, are used. Lining-papers serve as surfaces for oviposition and infusions provide the essential environment for nurturing. The types of oviposition surfaces and infusions used in laboratory rearing facilities or ovitraps influence oviposition preference, changing the breeding behavior of *Ae. aegypti* gravid mosquitoes. This study presents a comparative study on oviposition preferences for six different types of lining-papers and six different types of infusions in a bioassay cage. ANOVA analysis shows a significant effect on oviposition preference of different types of lining-papers that served as oviposition surfaces. The highest oviposition activity was observed for the *‘agri seed germination paper-75’* lining-paper when normal tap water infusion was used. Likewise, statistical analysis shows that when a *‘plain printing offset paper 80 GSM’* lining-paper was used, a highly statistically significant effect on oviposition preference is observed for different types of infusions used in the oviposition cups.

## 1. Introduction

*Aedes aegypti* (Linnaeus) female mosquitoes exhibit anthropophilic behavior and humans are their favorite supply of blood(McBride et al., 2014; Ponlawat & Harrington, 2005). It is one of the most prominent mosquito vectors responsible for transmitting viruses such as dengue, chikungunya, and Zika(Akiner et al., 2016). The magnitude of disease transmission by this vector raised serious health concerns. Since the start of 2023, five million confirmed cases and more than five thousand fatalities associated with dengue fever, reported across 80 countries in different regions of the World (Africa, Americas, South-East Asia, Western Pacific, and Eastern Mediterranean Regions)(WHO, 21 December 2023). A similar report was published by the European Centre for Disease Prevention and Control (ECDC) on 30 September 2023 for chikungunya. This report has revealed that the chikungunya virus infected about 0.44 million people between October 2022 and September 2023, with 350 deaths documented worldwide(ECDC, 2023).

*Ae. aegypti* thrives in urban and suburban environments due to the abundant presence of diverse oviposition sites that facilitate the deposition of eggs by this mosquito species(Dalpadado et al., 2022; Valença et al., 2013). For this reason, *Ae. aegypti* mosquitoes are capable of spreading vector-borne diseases rapidly.

The principal strategy for the management of vector-borne diseases of this characteristic involves effective vector control. However, controlling this mosquito species is extremely difficult because of its remarkable adaptive capacity and ability to deposit eggs in diverse locations, many of which are difficult to access(Carvalho & Moreira, 2017). In-depth understanding of the behavioral and breeding habits of *Ae. aegypti* is essential to control this vector by implementing an integrated pest management (IPM) approach where physical, biological, and chemical strategies can be used altogether(Abeyewickreme et al., 2018).

All mosquitoes undergo four unique stages throughout their life cycle: egg, larva, pupa, and adult (see **Fig. 1**). After consuming enough blood, gravid mosquitoes select a suitable oviposition site to lay their eggs(Day, 2016). *Ae. aegypti* gravid mosquitoes usually deposit their eggs on a surface around stagnant freshwater storage. Water is essential for mosquito breeding because mosquito eggs require some form of water to hatch. Mosquitoes use their hygro sensory system to detect humidity cues in order to explore their surroundings successfully in search of an appropriate oviposition site. After coming into direct contact with water, the female mosquitoes use their sensory and gustatory apparatus to detect the quality and texture of oviposition surfaces. A variety of suitable oviposition sites, such as old tires, clogged drains, canvas and plastic sheeting, and other stagnant water containers produced by human activities, are available for deposition of eggs(Day, 2016). The eggs that have been laid have the ability to live for extended periods. The eggs undergo hatching upon submerge in stagnant water. After hatching of eggs, the larvae feed on algae and other organic matters present in water(Merritt et al., 1992). Next, the larvae transform into pupae which also reside in water until they emerge as adult mosquitoes.

**Fig 1.**
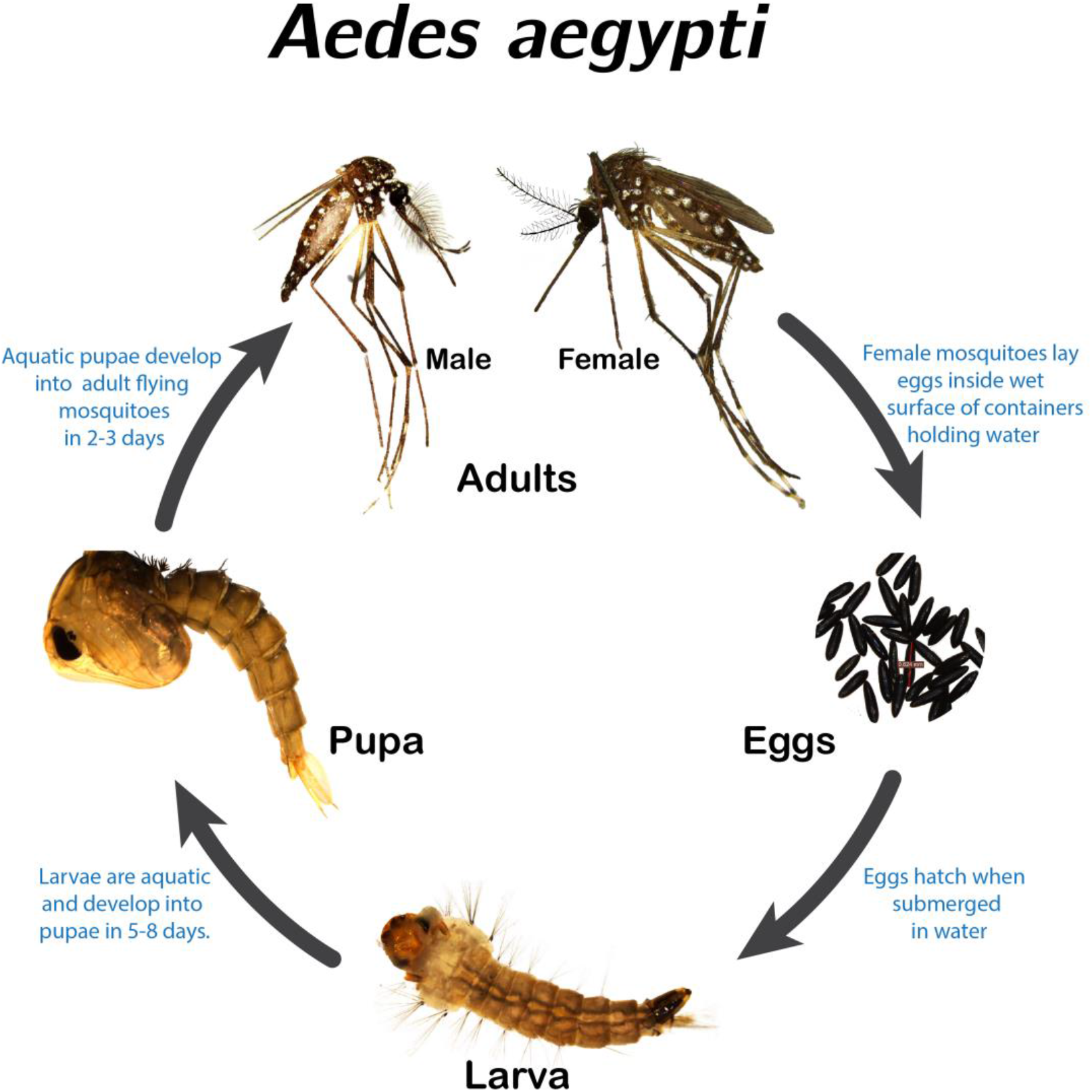
The entire life cycle of *Aedes aegypti:* egg, larva, pupa, and adult. This mosquito has an aquatic larva and a non-feeing pupa stage in its life cycle. The aquatic pupae are transformed into active flying adult mosquitoes at the final stage.

The egg stage is the most sensitive to environmental conditions in the first hours after being deposited(Sousa et al., 2023). For this reason, *Ae. aegypti* gravid mosquitoes show unique skip-oviposition behavioral pattern to ensure the maximum survival of the offsprings by distributing eggs in multiple suitable water containers(Colton et al., 2003). The selection of oviposition sites by a female mosquito is a conscious and analytical process, influenced by various physical and chemical factors present on an oviposition surface and in the water contents of a container(Mwingira et al., 2020). Water quality and contents in the water (i.e., presence of chemicals, microorganisms, etc.) of a container directly impact the aquatic larval and pupal stages of their life cycle till the adult stage. Therefore, in order to select an oviposition location, a gravid mosquito first assesses the nutritional quality of water as well as the likelihood of survival of their offsprings through visual and chemical cues. In addition, the availability of larval food and the presence of previous mosquito eggs or larvae influence the selection of oviposition sites(James et al., 2022; Montini & Fischer, 2024; Mwingira et al., 2020).

Eggs of *Ae. aegypti* absorb water in the early hours after egg deposition to facilitate embryogenesis. Serosal cuticle formation is completed after water absorption(Farnesi et al., 2015). For this reason, a proper moisture level of an oviposition surface is essential for freshly laid eggs in the first hour to form a darker dehydration-resistant protective cuticle(Bar-Zeev, 1967). As demonstrated in earlier research works, female mosquitoes exhibit a preference for a moist surface that is suitable for the survival and development of their eggs. In addition to the moisture level, gravid mosquitoes also consider the moisture absorbance capacity, temperature, surface texture, heat resistance capacity as well as the presence and absence of chemical substances and microorganisms in the damp surface of an egg-laying container(Bar-Zeev, 1967; Brouazin et al., 2022; Steinly et al., 1991).

In the bioassay cages and ovitraps (container breeding of *Aedes* spp.), various types of suitable lining-papers are used. The lining-papers can mimic the natural container inner surfaces of the preferred breeding sites of mosquitoes. However, there is a lack of comprehensive research on the effect of different types of inner surface lining-papers on oviposition site selection and oviposition preference(Bar-Zeev, 1967; Brouazin et al., 2022; Steinly et al., 1991). Furthermore, no comparative study was found that revealed the effect of different infusion and lining-paper combinations on oviposition preference.

Microorganisms are an essential food source for mosquito larvae. To mimic the natural breeding environment in the laboratory mosquito rearing facilities and ovitraps surveillance, occasionally a lab-made infusion containing organic substances is added to water-filled containers where mosquitoes deposit eggs. The infusions help to sustain the growth of microorganisms and serve as a food source for mosquito larvae. Artificially made infusions can be formulated from a variety of sources, including hay, grass, leaves, fruits, different plant extracts and even animal blood. Previous studies revealed that the type of infusions affects the oviposition behavior of gravid mosquitoes. For example, previous studies revealed that various semiochemicals and different plant organic infusions influenced oviposition preference and these infusions could be used to manipulate the skip-oviposition behavior (dispersing of eggs from a single batch across multiple oviposition sites instead of depositing all eggs in a single location) of gravid mosquitoes(Arbaoui & Chua, 2014; Mwingira et al., 2020; Obenauer et al., 2009).

However, there is a continuous search for more efficient and effective infusions that can be used in the containers of a bioassay cage and ovitraps to attract *Ae. aegypti* gravid mosquitoes for oviposition. Previous studies revealed that infusions comprising of larval rearing water and pre-existing larvae (conspecific and heterospecific) hold promise for a more sustainable, simple and cost-effective solution that is more specific for container breeders like *Ae. aegypti(Allan & Kline, 1995; Boullis et al., 2021; Tilak et al., 2005)*. However, limited information is available regarding these infusions containing larval-rearing water. Further investigation is required to reveal the influence and effectiveness of these infusions comprised of larval-rearing water on oviposition preference.

In this work, a comparative study on the effect of different types of oviposition surfaces and infusions on oviposition preference was carried out. First, the effect of six different types of lining-papers used as oviposition surfaces in combination with the normal tap wa*ter* infusion were investigated. Furthermore, the effect of six different types of infusions in combination with the *‘plain printing offset paper 80 GSM’* lining-paper was investigated. Our statistical analysis showed that different types of lining-papers and infusions significantly influence the oviposition preference of *Ae. aegypti* gravid mosquitoes.

## 2. Material and Methods

### 2.1. Laboratory Rearing of *Aedes aegypti* Mosquitoes

In this study, oviposition preference was investigated by counting eggs collected from a laboratory-maintained *Ae. aegypti* colony (see the workflow diagram in **Fig. 2**). In the laboratory, relative humidity, temperature, and light-dark cycle were consistently maintained at 60-80% RH, 27 °C (± 2 °C), and 12hL:12hD respectively. In the first step, mosquito eggs were collected from a laboratory-maintained *Ae. aegypti* colony (see **Fig. 3**) and transferred to a hatching tray containing water to keep the eggs wet in such a way that the lower part of the egg collecting strips were immersed in water. The newly hatched first instar larvae that emerged within six hours were collected using a dropper and placed into a rearing tray.

**Fig 2.**
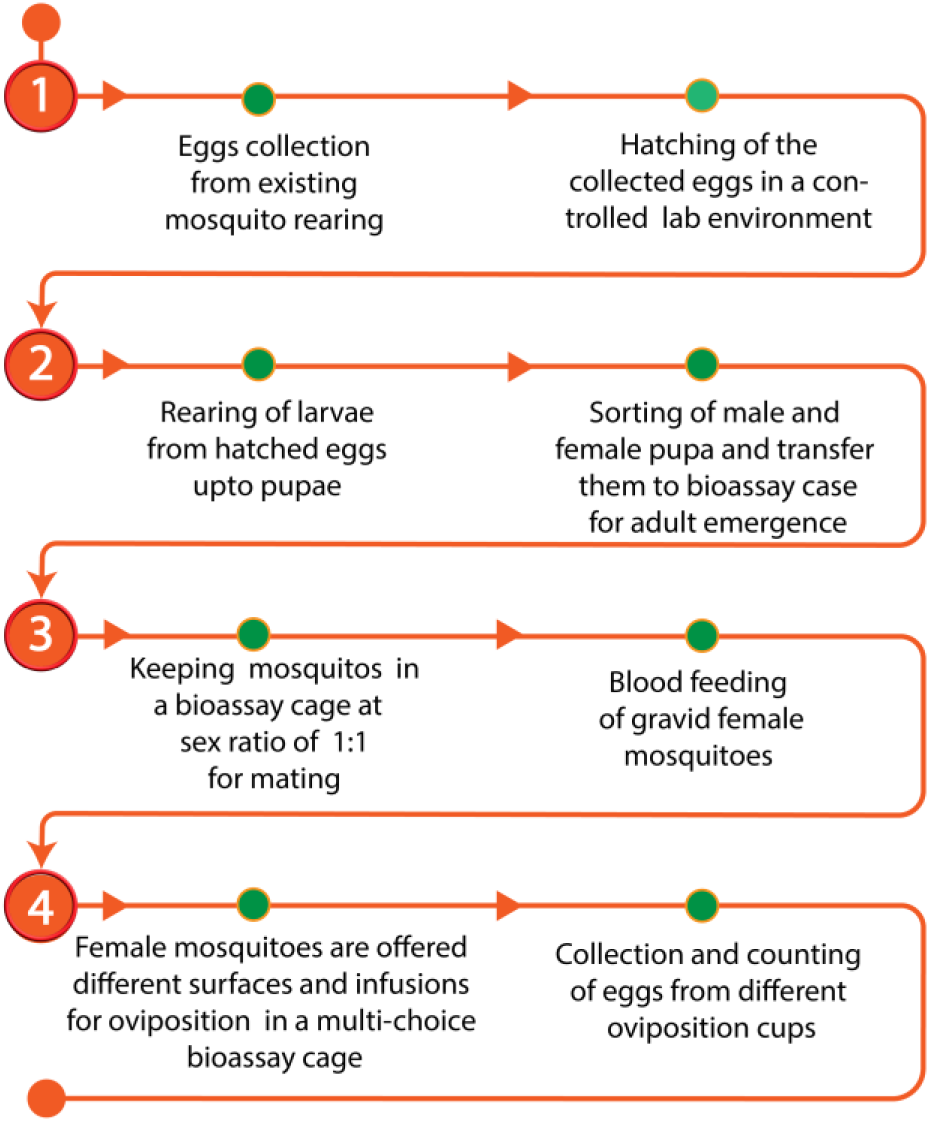
Steps for the experimental workflow. Our experiments started with collecting fresh eggs from an existing mosquito-rearing laboratory. To observe oviposition preference of gravid mosquitoes, eggs were collected from different oviposition cups containing different oviposition lining materials and infusions.

The larvae (see **Fig. 3 (b)**) were maintained in a larval rearing tray (40 cm × 27.30 cm) at a larval density of 1000 per tray containing 1000 ml water. Sufficient water was added from time to time to maintain the same water level during the larval rearing period. Daily 0.70 grams of fish feed (aquarium fish food Super Nova manufactured by Perfect Companion Group Co., Ltd. Thailand) was provided for 1000 larvae per tray. The rearing trays were cleaned periodically to remove excess waste materials using vacuum aspiration method (with the help of a vacuum pump). Once pupation was started, all the pupae were collected from the tray. The newly emerged pupae were collected, kept in plastic vials, and observed for adult eclosion.

A total of 40 adult mosquitoes (20 females and 20 males) at a 1:1 sex ratio was kept in a cage (35 cm × 28 cm × 20 cm size). Adult mosquitoes were provided with a daily supply of food using a fresh cotton ball soaked in a 10% glucose solution for the purpose of nourishing the adults. Furthermore, female mosquitoes were fed chicken blood by artificial membrane feeding (see **Fig. 3 (c, d)**), following a modification of the technique of Mourya *et al*.(Mourya *et al*., 2000). In this process, a blood bag containing anticoagulant (citrate phosphate dextrose) was used to collect chicken blood from the slaughterhouse of the local market. Small petri plates were filled with about 20 ml of blood (see **Fig. 3 (c, d)**) and then turned upside down on top of adult mosquito cages. Each petri plate was covered by a bag filled with warm water (around 37 °C) to keep the blood warm, where the female mosquitoes were permitted to blood feed for about 30–40 minutes. Female mosquitoes were allowed to consume blood for two consecutive days to ensure adequate blood feeding. After three days of being blood-fed, the females started laying eggs.

**Fig 3.**
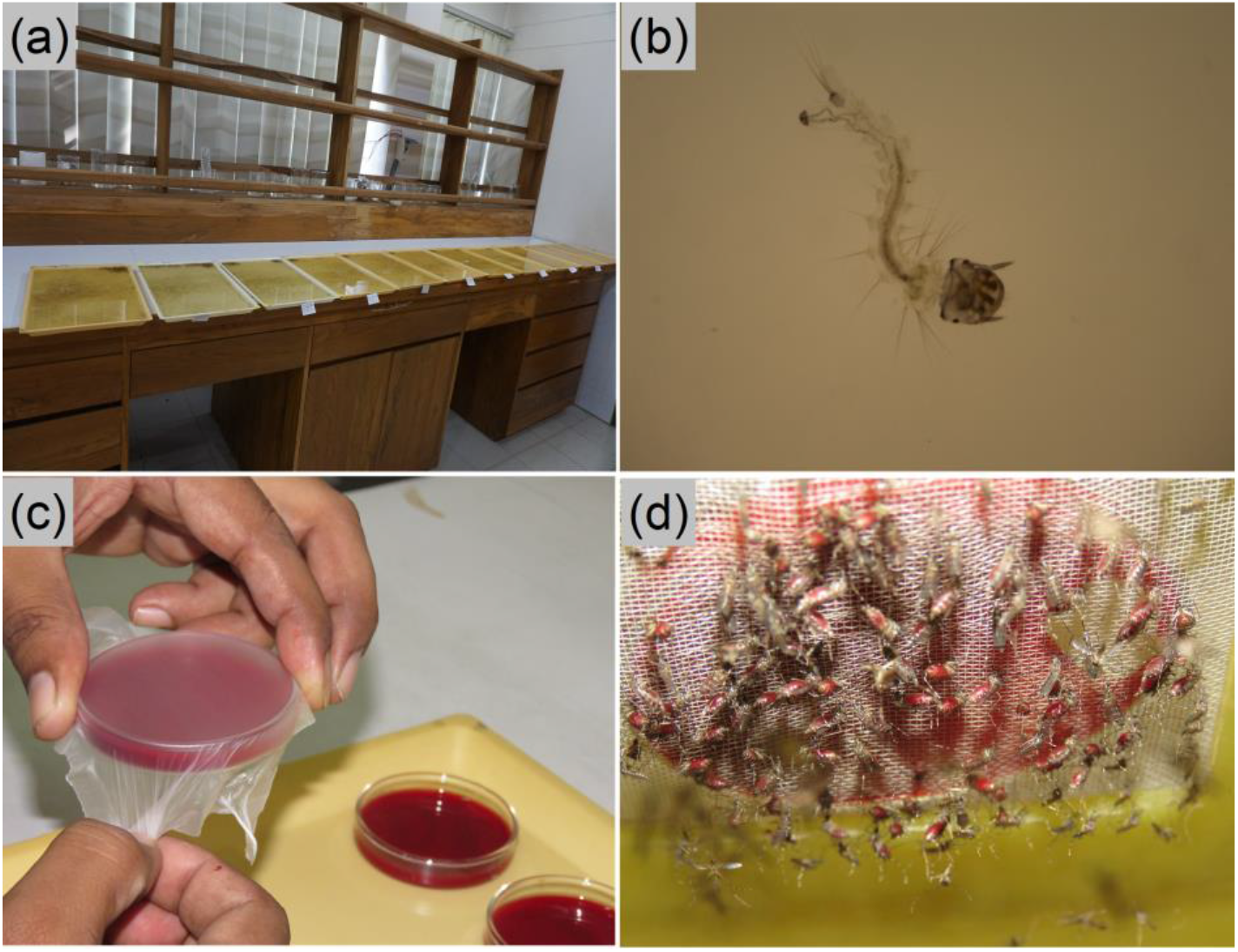
*Aedes* spp. rearing process (a) colony rearing room (b) larval stage (c) an artificial membrane blood-feeder setup preparation (d) appearance of female Aedes aegypti mosquitoes that have engorged.

To investigate oviposition preference, a bioassay cage for a multi-choice test was used (see **Fig. 4**). Six oviposition cups made of clear plastic were placed in the bioassay cage. Inner surfaces of the cups were covered using different lining papers (see **Table 1**) which were functioned as oviposition surfaces.

**Fig 4.**
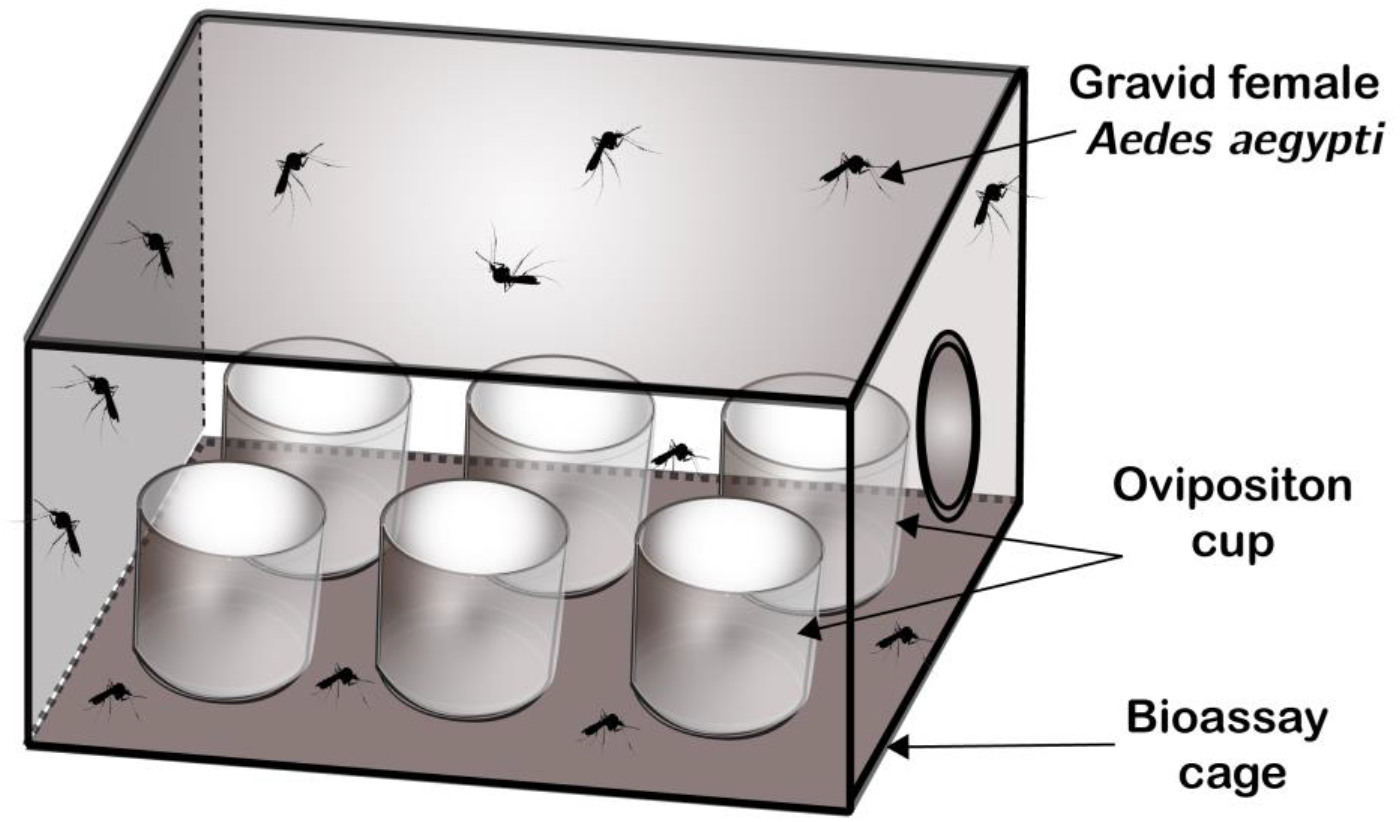
Schematic diagram of a bioassay cage for a multi-choice test for investigating oviposition preference. To study the effect of different lining-papers that functioned as oviposition surfaces, six types of papers were used as inner surface lining on the oviposition cups (breeding container) and ‘*normal tap water’* infusion was added. Similarly, to study the effect of different infusions, six types of infusions were added for each oviposition cup and ‘*plain printing offset paper 80 GSM*’ was used as a lining-paper that functioned as oviposition surface.

**Table 1.** Properties of different types of lining-papers and their code name used in this experiment.

| Lining-paper<br>(code name) | Type of paper and<br>manufacturer | Grams per square meter<br>(GSM) | Thickness<br>($\mu\text{m}$) | Texture | Average surface<br>Roughness, Ra<br>($\mu\text{m}$) | Color |
| --- | --- | --- | --- | --- | --- | --- |
| OS-A | Plain printing offset<br>paper 80 GSM<br>(Bashundhara Paper<br>Mills Limited,<br>Bangladesh) | 80 | 92.5±2.52 | plain | 2.91 | white |
| OS-B | Qualitative filter paper<br>(Whatman<sup>®</sup>) | 87 | 180±0.95 | plain | 9.08 | white |
| OS-C | Agri seed germination<br>paper KK030 (Kalpkala<br>Paper & Pulp Paper<br>Industries, India) | 160 | 374.4±<br>51.56 | crepe | 60.62 | white |
| OS-D | Agri seed germination<br>paper KK029/85<br>(Kalpkala Paper & Pulp<br>Paper Industries, India) | 84 | 176±56.71 | crepe | 43.46 | white |
| OS-E | Agri seed germination<br>paper-125 (Kalpkala<br>Paper & Pulp Paper<br>Industries, India) | 130 | 262.88±<br>48.58 | crepe | 97.90 | light<br>shade<br>of brown |
| OS-F | Agri seed germination<br>paper-75 (Kalpkala<br>Paper & Pulp Paper<br>Industries, India) | 79 | 156.29±<br>58.45 | crepe | 57.21 | warm and<br>earthy<br>shade of<br>brown |

We obtained those six types of lining papers from different manufacturers for our study and identified them by their respective commercial names. Details of lining papers with their respective code name used in this article is presented in **Table 1**. Images of the surfaces of the lining papers are also shown in **Fig. 5**. In addition, surface profiles of the lining papers were investigated by a stylus profilometer (Dektak 150 Surface Profiler, Veeco Instruments Inc.) as shown in **Fig. 6**. A lateral scan of 5000 μm was performed using a stylus of radius 12.5 μm (force 1.0 mg) for each lining paper. From the surface profiles, average roughness *R*_*a*_ (see **Table 1)** of each lining paper were calculated using the following equation:

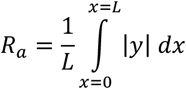

Here, *y* is the surface height and R_a_ is the arithmetic average deviation from the mean line within the lateral scan length (L) in the *x* direction.

**Table 2** presents the six different types of infusions with their code name used in this article. Please note that the *normal tap water (IN-B)* is also termed as infusion in this article.

**Fig 5.**
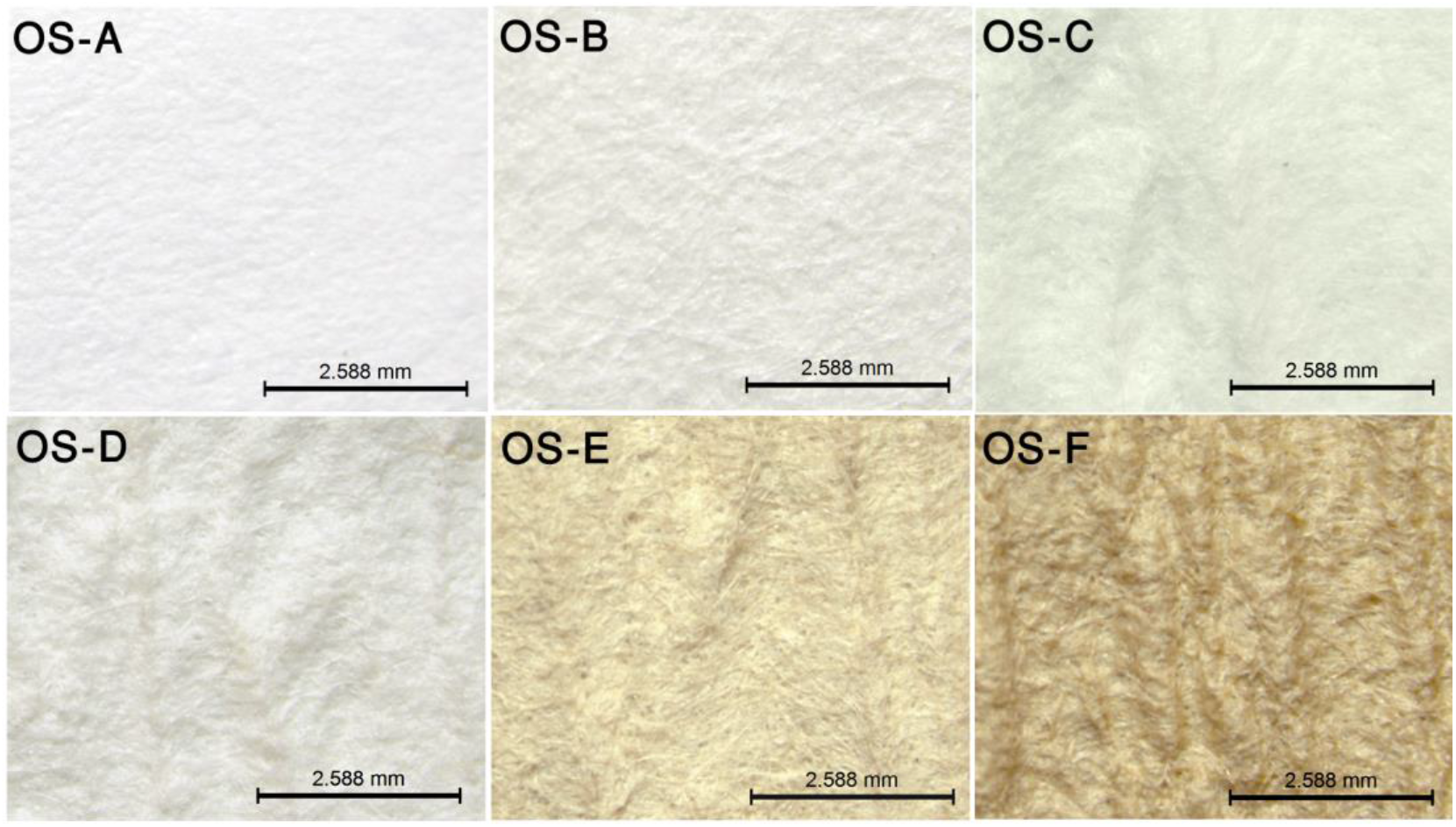
Images of different lining paper surfaces: OS-A (plain printing offset paper 80 GSM); OS-B (qualitative filter paper); OS-C (agri seed germination paper KK030); OS-D (agri seed germination paper KK029/85); OS-E (agri seed germination paper-125); OS-F (agri seed germination paper-75).

**Fig 6.**
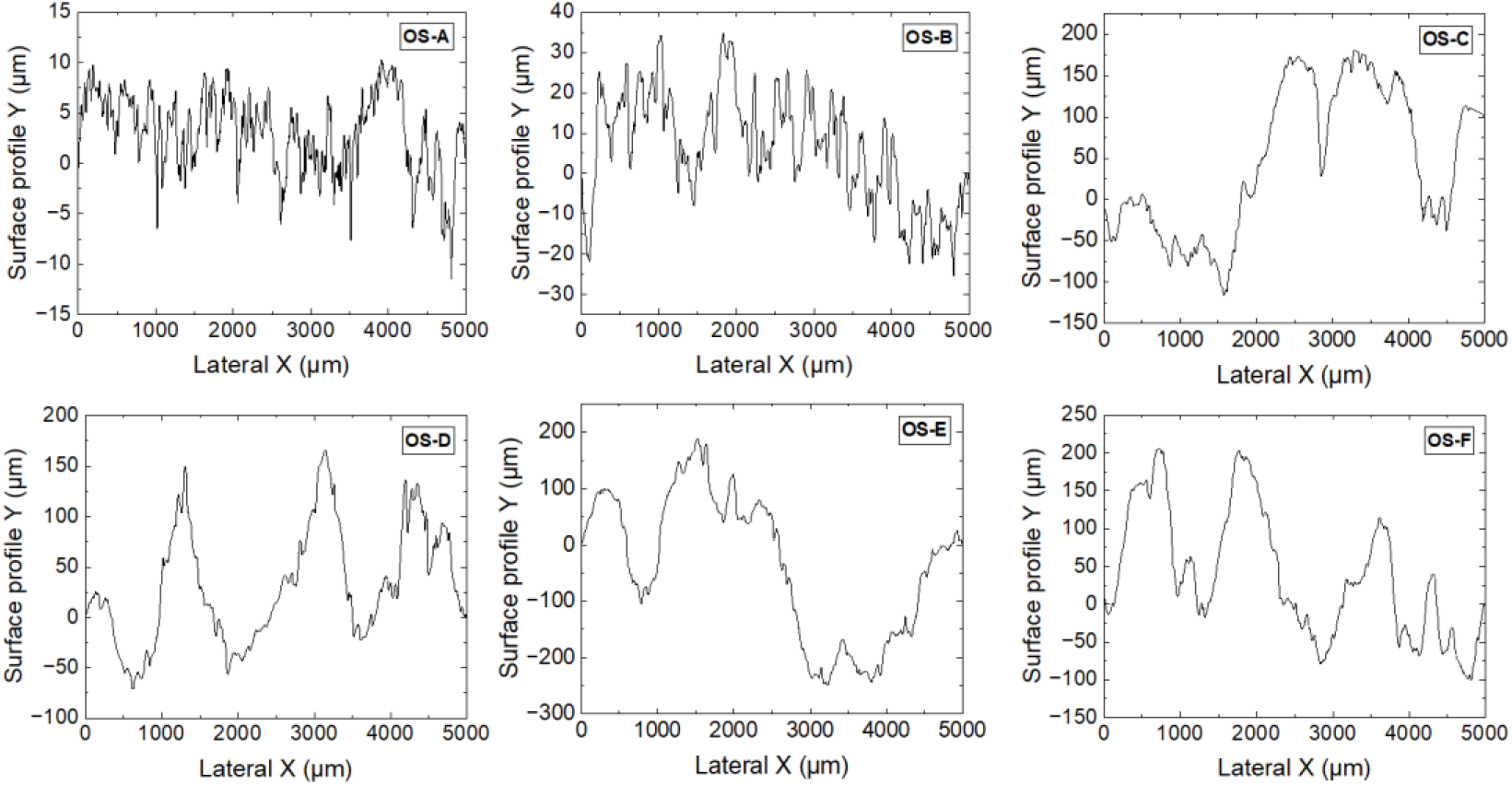
Surface profile of different types of lining papers: OS-A (plain printing offset paper 80 GSM); OS-B (qualitative filter paper); OS-C (agri seed germination paper KK030); OS-D (agri seed germination paper KK029/85); OS-E (agri seed germination paper-125); OS-F (agri seed germination paper-75).

As *Ae. aegypti* females do not lay eggs in the absence of a wet surface(Bar-Zeev, 1967), therefore oviposition cups with the *normal tap water (IN-B)* and the *plain printing offset paper 80 GSM (OS-A)* lining were considered as control in this study. In addition, the *OS-A* lining paper and *IN-B* infusion were used for regular colony maintenance of *Ae. aegypti*.

This study was divided into two separate experiments. In the first experiment, after blood feeding the gravid mosquitoes were given a choice of six oviposition cups to lay eggs where different types of lining-papers along with *normal tap water (IN-B)* were used in each oviposition cup (see **Table 3**). Similarly, in the second experiment, *Ae. aegypti* gravid mosquitoes were given a choice to lay eggs between six oviposition cups where different infusions were added in each oviposition cup along with plain printing offset paper 80 GSM (OS-A) (see **Table 3**). Each oviposition cup is provided with an equal amount of infusion. After three days of oviposition, each lining-paper was collected from the oviposition cups to count the number of eggs. Each experiment was repeated three times.

### 2.2. Data Analysis

The effect of different oviposition surfaces and water composition was expressed by the oviposition activity index (OAI) as described by Kramer and Mulla(Kramer & Mulla, 1979). The oviposition activity index (OAI) was calculated by the following formula:

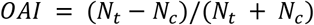

where, N_t_ represents the number of eggs observed on the lining-material (oviposition surface) of the oviposition cups, and N_c_ represents the number of eggs observed on the lining-material (oviposition surface) of the control oviposition cup.

The oviposition activity index (OAI) is bounded within the interval of +1 to −1, where 0 indicates no response. A positive value indicates that the infusion or lining material of an oviposition cup functions as an attractant, whereas a negative value indicates that the infusion or surface has a deterrent effect.

One-way analysis of variance (ANOVA) was used to compare the influence of the six different types of oviposition surfaces (lining-papers) as well as six different types of infusions. The Fisher’s Least Significant Difference (LSD) test was used to compare the different oviposition surfaces as well as different infusions. In this context, P ≤ 0.05 was referred to as statistically significant and P ≤ 0.001 was referred to as statistically highly significant. The statistical analyses were carried out by OriginPro ® software.

## 3. Results

### 3.1. Oviposition Preference for Different Types of Paper

In the experiment of oviposition preference for different lining-paper surfaces, the *plain printing offset paper 80 GSM (OS-A)* lining-paper attached to the inner surface of the oviposition cups was considered as control (see **Table 1**).

Overall, the mean number of eggs laid in the oviposition cups by the gravid mosquitoes varied depending on the types of lining-papers functioned as oviposition surface (see **Fig. 7 (a)**). Statistical analysis showed statistically significant (F = 4.852; df = 5, 12; P = 0.01) evidence of correlation of oviposition preference with different lining-papers (see **Fig. 7 (b)**). The highest amount of oviposition activity (see **Fig. 8**) was observed in the case of the *agri seed germination paper-75 (OS-F)*. According to mean comparison conducted in pairs using the Fisher’s Least Significant Difference (LSD) test, only the effect of *agri seed germination paper-75 (OS-F)* paper lining was significantly different from other five types of lining-paper.

**Fig 7.**
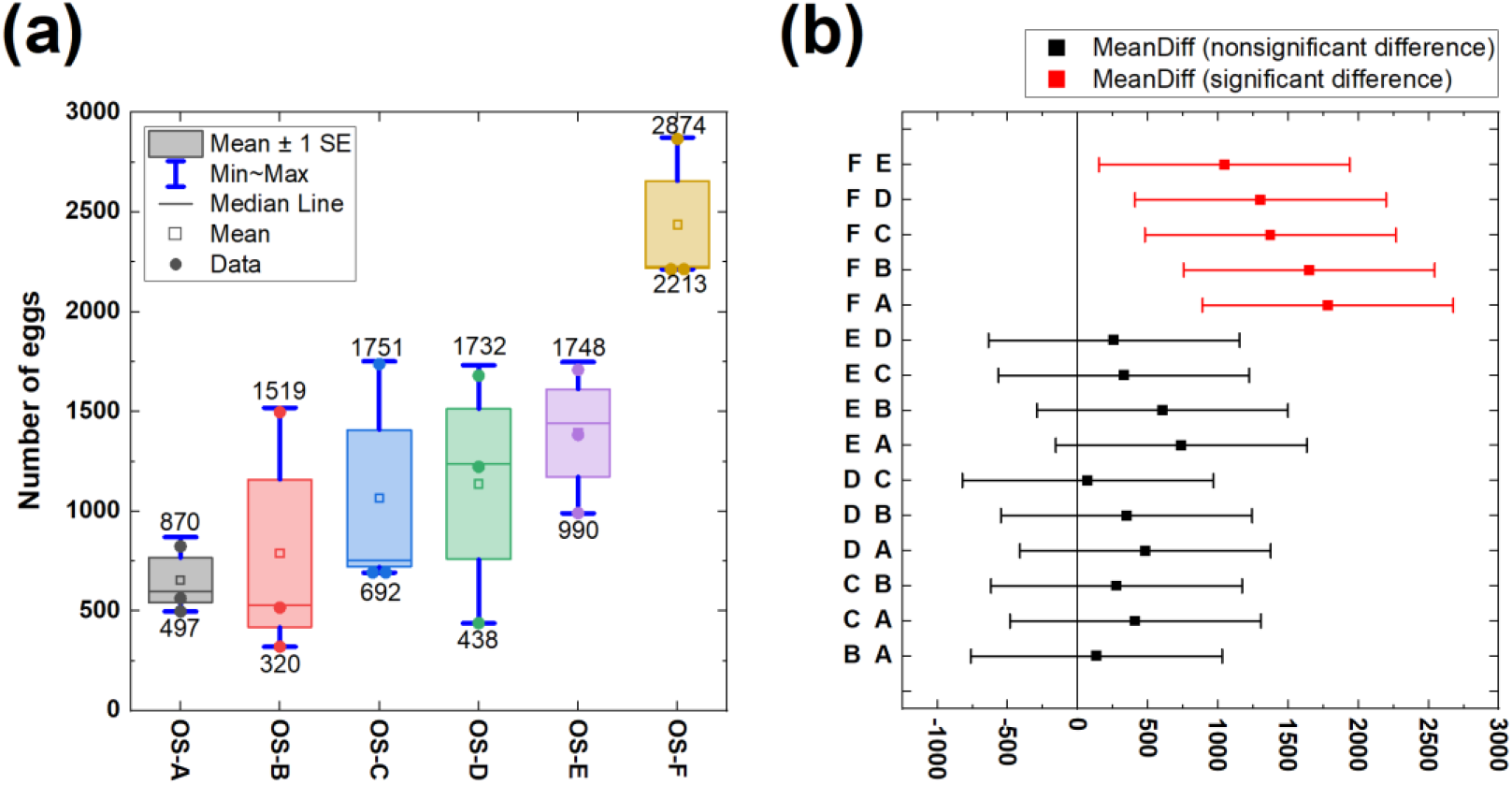
Oviposition preference of *Aedes aegypti* gravid mosquitoes in response to the type of lining-paper surface of an oviposition container (a) effect of six types of lining-papers functioned as an oviposition surface (see Table 1) (b) means comparison using Fisher’s Test. Here, A (plain printing offset paper 80 GSM), B (qualitative filter paper), C (agri seed germination paper KK030), D (agri seed germination paper KK029/85), E (agri seed germination paper-125), F (agri seed germination paper-75).

**Fig 8.**
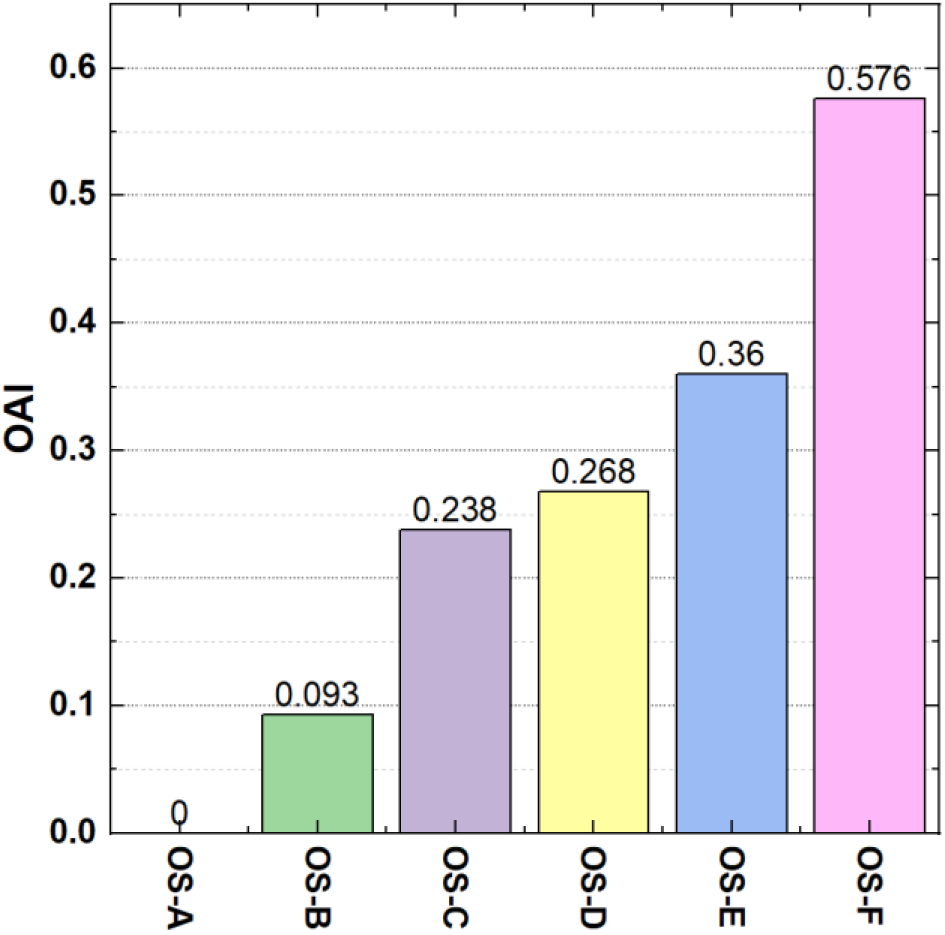
Oviposition activity index (OAI) for a multi-choice test of *Aedes aegypti* gravid mosquitoes corresponding to different types of lining-papers (see Table 1) functioned as oviposition surfaces surrounding inner surface of an oviposition cup. Here, OS-A (plain printing offset paper 80 GSM), OS-B (qualitative filter paper), OS-C (agri seed germination paper KK030), OS-D (agri seed germination paper KK029/85), OS-E (agri seed germination paper-125), OS-F (agri seed germination paper-75).

In the case of *plain printing offset paper 80 GSM (OS-A)*, the value of the Oviposition Activity Index (OAI) was zero. For the rest of the five different oviposition lining-papers, the values of the Oviposition activity index (OAI) appeared between 0 and 0.576 (see **Fig. 8**).

### 3.2. Oviposition Preference for Different Types of Infusions

In the second experiment, the oviposition preference of *Ae. aegypti* gravid mosquitoes for different infusions was investigated. In this case, the *plain printing offset paper 80 GSM (OS-A)* was used as lining-paper which functioned as an oviposition surface attached to the inner surface of oviposition cups (see **Table 2**).

**Table 2.**
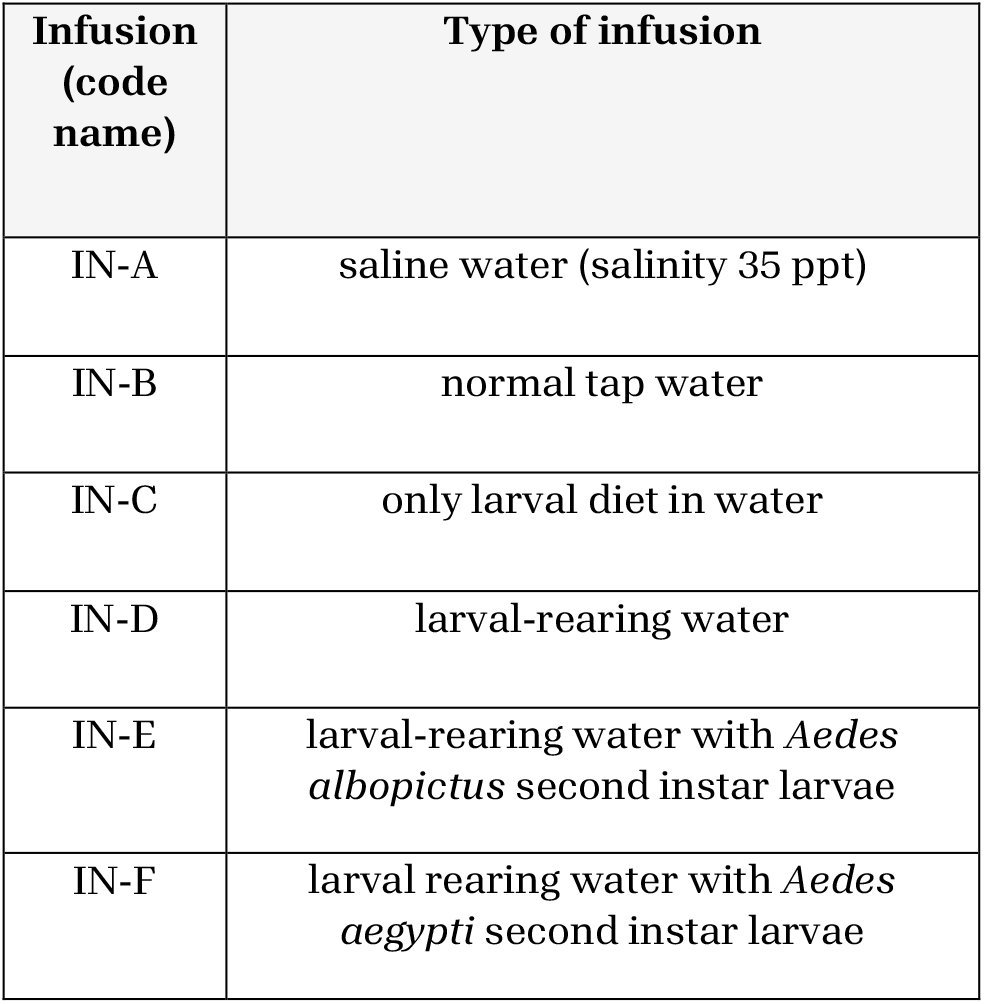
Different types of infusions and their code names were used in this experiment.

| Infusion (code name) | Type of infusion |
| --- | --- |
| IN-A | saline water (salinity 35 ppt) |
| IN-B | normal tap water |
| IN-C | only larval diet in water |
| IN-D | larval-rearing water |
| IN-E | larval-rearing water with *Aedes albopictus* second instar larvae |
| IN-F | larval rearing water with *Aedes aegypti* second instar larvae |

**Table 3.** Combination of different types of infusions and lining-papers used to observe the effect on oviposition preference.

| Experiment 1:<br>Effect of different<br>oviposition surfaces |  | Experiment 2:<br>Effect of different<br>infusions |  |
| --- | --- | --- | --- |
| Lining-paper (code name) | Type of infusion (code name) | Infusion (code name) | Lining-paper (code name) |
| OS-A | IN-A | IN-A | OS-A |
| OS-B | IN-A | IN-B | OS-A |
| OS-C | IN-A | IN-C | OS-A |
| OS-D | IN-A | IN-D | OS-A |
| OS-E | IN-A | IN-E | OS-A |
| OS-F | IN-A | IN-F | OS-A |

Overall, the mean number of eggs laid in oviposition cups by the gravid mosquitoes varied depending on the types of infusions added in the infusion cups (see **Fig. 9 (a)**). Statistical analysis showed a statistically highly significant (F = 8.629; df = 5, 12; P = 0.001) correlation of oviposition preference with different water compositions (infusions) used in the oviposition cups. The mean comparison in pairs using Fisher’s Least Significant Difference (LSD) test (see **Fig. 9 (b)**) showed that the infusion containing *larval rearing water with Ae. aegypti larvae (IN-F)* were significantly different when compared with the *saline water (IN-A), normal tap water (IN-B)*, or *only larval diet in water (IN-C)*. In the case of *larval rearing water with Ae. albopictus larvae (IN-E)*, the pairwise Fisher’s LSD mean comparison was significantly different from *saline water (IN-A)* and *normal tap water (IN-B)* (see **Fig. 9 (b)**). According to Fisher’s LSD mean comparison test, no statistically significant effect (see **Fig. 9 (b)**) was observed between the *larval rearing water with Ae. albopictus larvae (IN-E)* and *larval rearing water with Aedes aegypti larvae (IN-F)*.

**Fig 9.**
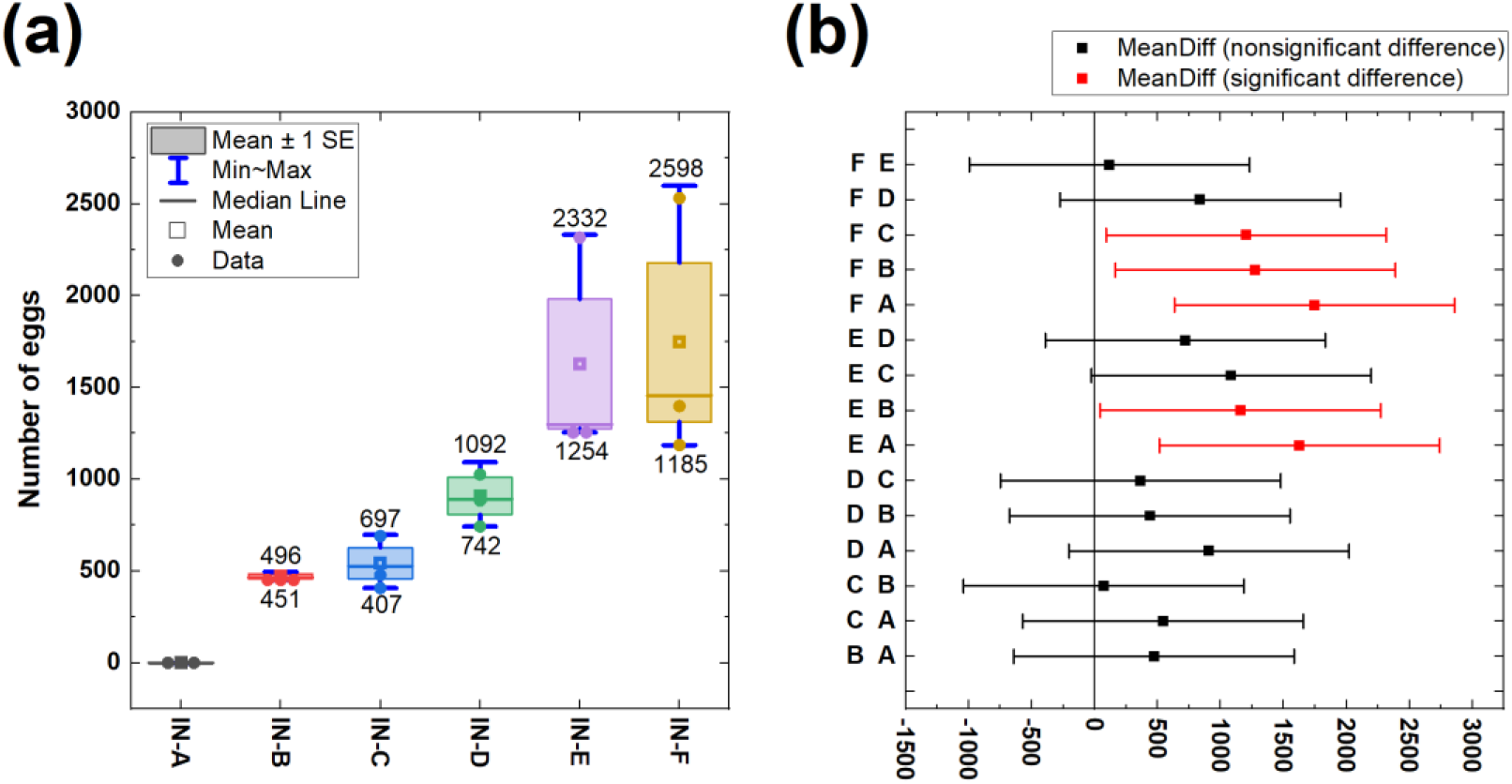
Oviposition preference of *Aedes aegypti* gravid mosquitoes corresponding to different infusions (a) effect of six types of infusions in oviposition cups (see Table 2) (b) means comparison using Fisher’s Test. Here, A (saline water), B (normal tap water), C (only larval diet in water), D (Aedes aegypti larval-rearing water), E (larval-rearing water with *Aedes albopictus* larvae), F (larval rearing water with *Aedes aegypti* larvae).

In this case, the *normal tap water (IN-B)* for laboratory colony maintenance was considered as a control, where the value of the Oviposition Activity Index (OAI) was zero. For the rest of the five different infusions used in the oviposition cups, the values of the Oviposition activity index (OAI) appeared between −1 and 0.5754 (see **Fig. 10**).

**Fig 10.**
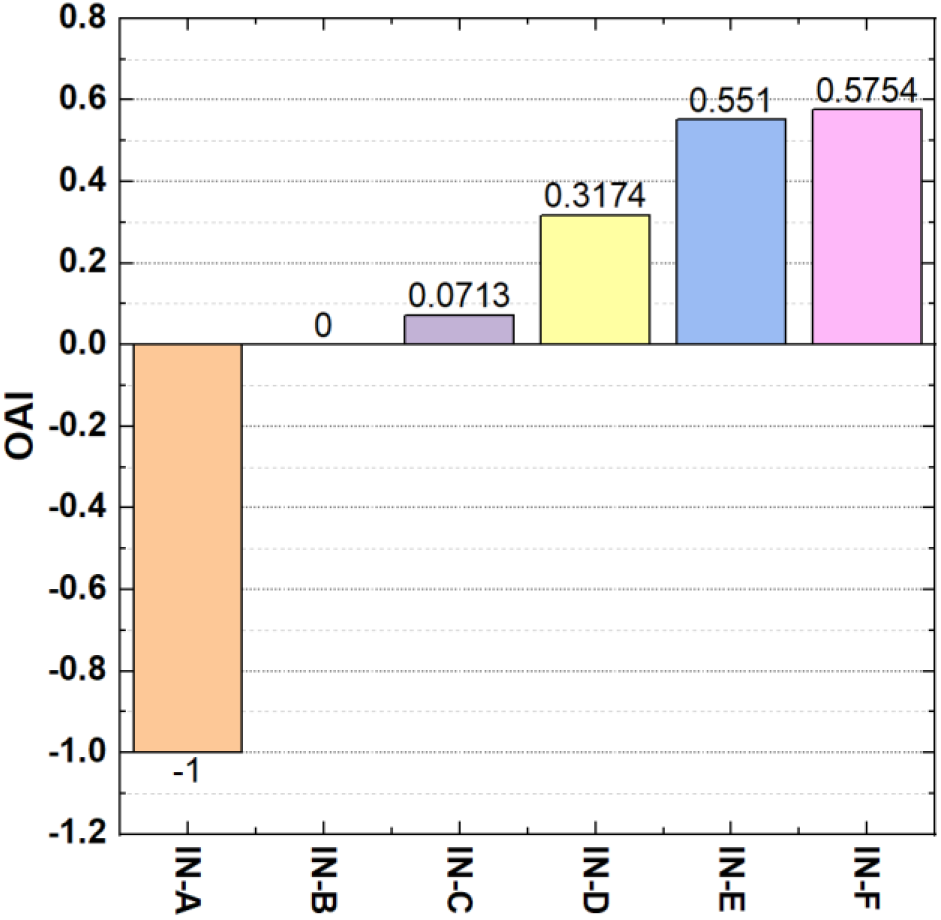
Oviposition activity index (OAI) for a multi-choice test of *Aedes aegypti* gravid mosquitoes corresponding to different infusions (see Table 2) used in the oviposition cups located inside a bioassay cage. Here, IN-A (saline water), IN-B (normal tap water), IN-C (only larval diet in water), IN-D (*Aedes aegypti* larval-rearing water), IN-E (larval-rearing water with *Aedes albopictus* larvae), IN-F (larval rearing water with *Aedes aegypti* larvae).

The highest oviposition activity was seen in the case of conspecific larval rearing water with their second instar larval stage (IN-F) used in the oviposition cup (see Fig. 10). All types of infusions in the oviposition cups functioned as attractant except the *saline water (IN-A)*. The application of *saline water (IN-A)* infusion has functioned negatively to oviposition preference.

## 4. Discussion

### 4.1. Effect of Different Oviposition Surfaces

*Aedes aegypti* gravid mosquitoes prefer to lay their eggs on rough and moist surfaces of the oviposition sites(Brouazin et al., 2022; Friuli et al., 2022; Madeira et al., 2002; O’Gower, 1957). It was reported that *Aedes* spp. expresses their preference to an oviposition surface with a moisture level similar to 70% of the soil SMC (saturation moisture content) to ensure the embryogenesis of eggs(Knight & Baker, 1962).

In our experiment, we used different types of lining-papers that served as oviposition surfaces. The lining-papers attached to the inner surface of the oviposition cups/containers mimic the natural surface in laboratory conditions. As different types of lining-papers have different water retention capacities as well as wet strengths, it was anticipated that the various types of lining-papers would have an impact on the oviposition preference. Our results also showed that depending on the types of lining-papers, *Ae. aegypti* gravid mosquitoes showed statistically significant different oviposition preferences. In addition, our results also showed that seed germination papers of different variants (see **Fig. 7** and **Fig. 8**) are more suitable as an oviposition surface due to higher oviposition activity in comparison with the *plain printing offset paper 80 GSM (OS-A)* or *qualitative filter paper (OS-B)*. The higher oviposition activity index was attributed to the higher water retention capacity and surface roughness of the different variants of seed germination papers. The highest oviposition activity was observed in the case of *agri seed germination paper-75 (OS-F)*. Further research work is required to explore the exact reason for higher OAI in the case of *OS-F* paper.

### 4.2. Effect of Different Infusions

As the availability of food is essential for the survival of larvae, female mosquitoes were expected to choose oviposition sites containing rich organic matter to ensure food for their offspring(Albeny-Simões et al., 2014; Kroth et al., 2019). In contrast to the findings of Brouazin *et al*.*(Brouazin et al*., *2022)*, our results evidently demonstrated that gravid mosquitoes prefer water containing a larval meal over normal tape water (see **Fig. 9** and **Fig. 10**). This finding can be explained by the fact that the water in which mosquito larvae and pupae developed previously contained larval-produced excretory products and microbiota-derived metabolites that provided oviposition cues to attract gravid mosquitoes(Brouazin et al., 2022; Mosquera et al., 2023). Those cues simulated naturally occurring oviposition pheromones specifically to help gravid mosquitoes in recognizing suitable oviposition sites for the development of newly hatched larvae(Albeny-Simões et al., 2014; Gunathilaka et al., 2018; Kalpage & Brust, 1973). Therefore, the presence of conspecific larvae in the infusions can be an indicator of a superior oviposition site. Likewise, the microbiota of larval rearing water also plays a vital role in generating an odorant that attracts gravid mosquitoes to the oviposition site(Arbaoui & Chua, 2014; Melo et al., 2020). For example, previous research revealed that *Aedes togoi* and *Ae. atropalpus* female mosquitoes show oviposition preference for the water (infusion) containing conspecific larvae reared in a sterile condition or water (infusion) in which fourth instar larvae were kept for 48 hours. Moreover, their oviposition preference also depends on the density of the larvae in order to ensure enough resources for preadult development(Kalpage & Brust, 1973; Trimble & Wellington, 1980).

Our investigation revealed that, *Ae. aegypti* gravid mosquitoes display a significantly positive oviposition preference to conspecific larval rearing water (*IN-E* and *IN-F*) over all the other infusions used in our experiment (see **Fig. 9** and **Fig. 10**). This finding also supports the previous study carried out by Allan & Kline where they observed similar oviposition responses between conspecific larval water and normal water(Allan & Kline, 1998). However, they used larval rearing water stored at frozen temperature (−20 °C for up to 3 weeks) whereas in this experiment we used larval rearing water stored at the temperature of 27±1 °C for 24 hours before the experiment. The findings of our experiment were also consistent with several previous research works involving larval water(Brouazin et al., 2022; Kalpage & Brust, 1973; Serpa et al., 2008; Wong et al., 2011). The oviposition preference pheromone associated with the presence of larval stage was elucidated by investigating behavioral and antennal responses of *Ae. aegypti* female mosquitoes to chemical cues from conspecific larvae(Boullis et al., 2021; Faierstein et al., 2019; Gonzalez et al., 2014). For example, oviposition pheromone for *Ae. aegypti* and *Ae. albopictus* n-heneicosane (C21) was detected from the cuticle of their larval stage(Gonzalez et al., 2014). Carboxylic acid compounds such as 9-Octadecenoic acid (Z)-, methyl ester, dodecanoic acid, and tetradecanoic acid isomers extracted from larva were also observed to play a key role in the localization of oviposition sites by *Ae. aegypti* gravid mosquitoes(Wang et al., 2019).

In addition, salinity detection of the oviposition sites is also very important for the survival of *Ae. aegypti* larvae as contamination of water with seawater (as little as 12.5%) becomes lethal for their larvae(Ponnusamy et al., 2008). In order to ensure larval survival, the oviposition preference of gravid mosquitoes decreases as salinity increases(Clark et al., 2004; Gunathilaka et al., 2018). To detect lethal concentrations of saline water for their larvae, *Ae. aegypti* mosquitoes use specific neurons in their legs and mouthparts(Matthews et al., 2019). For this reason, in our experiment it was observed that the oviposition cup, containing simulated seawater (*IN-A)* with water salinity of about 35 ppt, repeals *Ae. aegypti* gravid mosquitoes from laying eggs (see **Fig. 9** and **Fig. 10**). However, some *Aedes* species, for example *Ae. vigilax*, may prefer to oviposit in salt water(Knight et al., 2012).

It is evident from our study that larval rearing water with a conspecific larval stage (*IN-E* and *IN-F*) offers more cues to aid *Ae. aegypti* gravid mosquitoes in selecting their oviposition site compared to normal tap water (*IN-B*) or water containing only a larval diet (*IN-C*).

The findings from our oviposition preference study showed that both the types of lining-papers along with infusions used in oviposition cups have a significant effect on the oviposition preference for Ae. aegypti gravid mosquitoes. While preliminary, our study suggests that the *agri seed germination paper-75 (OS-F)* lining-paper in combination with the *larval rearing water with Ae. aegypti larvae (IN-F)* infusion could be used to design and develop a better oviposition cup that can be used in a bioassay cage or ovitrap(Allan & Kline, 1995; Leite Alves et al., 2018; Mackay et al., 2013; Obenauer et al., 2009; Parker, 2020; Ritchie et al., 2003; Shu & Shelomi, 2021).

## 5. Conclusion

In-depth understanding of the behavioral and breeding habits of *Aedes aegypti* is essential to control this vector by implementing an integrated pest management (IPM) approach where physical, biological, and chemical strategies can be used altogether. To study the behavioral and breeding habits of *Ae. aegypti*, a mimic of the natural breeding sites is usually built in a laboratory where highly efficient, species-specific container breeding mosquito population can be maintained as well as the effect of varying controlled environmental parameters can be observed (Day, 2016). In laboratory rearing of *Ae. aegypti* mosquitoes, oviposition preference of *Ae. aegypti* gravid mosquitoes varies depending on the types of lining materials that function as oviposition surfaces attached to the inner surface of oviposition cups or containers. In addition, the infusions used in oviposition cups show a significant effect on oviposition preference. Understanding *Ae. aegypti* mosquito breeding behavior which is influenced by different types of lining materials and infusions is essential to design and develop a better laboratory-rearing facility or ovitrap. This work presents a comparison study of the effect on the oviposition preference for six different types of lining-papers served as oviposition surfaces as well as six different types of infusions added in oviposition cups in a bioassay cage. It was observed that both the types of lining-papers along with infusions used in oviposition cups have a significant effect on the oviposition preference for *Ae. aegypti* gravid mosquitoes. Among the different types of lining-papers that functioned as oviposition surfaces, the highest oviposition preference was observed for the *agri seed germination paper-75 (OS-F)* in combination with the *normal tap water (IN-B)* infusion. Furthermore, six different types of infusions in the oviposition cups along with the *plain printing offset paper 80 GSM (OS-A)* lining-paper were used. In comparison with the *normal tap water (IN-B)* infusion, the *saline water (IN-A)* infusion showed a negative effect on the oviposition preference. The *larval rearing water with Aedes aegypti larvae (IN-F)* infusion showed the highest oviposition preference. Our study suggests that the *agri seed germination paper-75 (OS-F)* lining-paper in combination with the *larval rearing water with Aedes aegypti larvae (IN-F)* infusion could be used to design and develop a better oviposition cup that can be used in a bioassay cage or ovitrap. However, further research work is required to explore the key factors, such as water retention capacity and texture pattern of the oviposition surface of lining-papers that influence the oviposition preference of the gravid mosquitoes to a particular type of seed germination paper.

## List of Abbreviations

Ae.: *Aedes*
GSM: grams per square meter
ECDC: European Centre for Disease Prevention and Control
IPM: Integrated pest management
RH: Relative humidity
OAI: oviposition activity index
LSD: Least Significant Difference

## CRediT authorship contribution statement

### Mahfuza Momen

Conceptualization, Data curation, Formal Analysis, Funding acquisition, Investigation, Methodology, Project administration, Resources, Software, Supervision, Validation, Visualization, Writing original draft, Writing review & editing. **Kajla Sheheli:** Funding acquisition, Writing review & editing. **Md. Aftab Hossain:** Writing review & editing. **Ananna Ghosh:** Investigation, Writing-review & editing. **Md. Forhad Hossain:** Writing-review & editing.

## Data availability

The experimental data will be provided upon request.

